# Across-lineage prediction of within-KO sequence diversity by temperature and pH differs among subcellular compartments in hydrothermal spring communities

**DOI:** 10.1101/2025.06.12.659262

**Authors:** Juan Rivas-Santisteban, Jaime Alcorta, Beatriz Díez, Javier Tamames, Carlos Pedrós-Alió

## Abstract

A general understanding of how extreme temperature and pH might universally shape the evolution of protein variants is lacking. Importantly, while pH is differentially regulated in microbial compartments, temperature is not. We hypothesise that these variables may partly explain the within-genome variability in protein rates of single-celled organisms, since genes encode proteins allocated to different compartments. To test this hypothesis, we examine the number of unique sequence contributors assigned to a KEGG across species as a coarse proxy for accumulated evolutionary divergence. Under a naive null expectation, orthologs of equal abundance should be represented by a similar number of variants. We examine sequence diversity in 17 metagenomes from El Tatio geothermal field (Chile), spanning temperatures of 45-62°C, and pH values of 7.2-9.3. At equal abundance, warmer temperature is weakly associated to more ortholog variants, while alkaline pH is associated to fewer (in the ranges examined). The inclusion of these variables in the model robustly improved the prediction of the community-level within-KO unique sequence contributors from metagenomic abundance. KO variant diversity in the periplasm was only significantly affected by pH increases, providing partial support to the compartment hypothesis.

## Introduction

(1) How does the overall number of sequence variants scale with the metagenomic abundance of a KEGG annotation? (2) Which factors influence this prediction over evolutionary timescales? While question (1) is deceptively straightforward (the more abundant a KEGG is across taxa, the higher number of sequence variants will contribute to its abundance), the resolution of question (2) may further elucidate the causes underlying the observed functional diversity. The number of distinct sequences assigned to a KEGG orthology ID is not a direct count of evolutionary events, but it provides a coarse proxy for the accumulated evolutionary history of that function across taxa [1–5], as it recalls mutation [6], duplication [7], indels [8], length variation [9], horizontal transfer [10, 11], gene loss [12], genome reduction [13], tandem and antiparallel gene overlapping [14], change in GC content [15], among others. Thus, knowing which factors make the by-KEGG sequence diversity grow or diminish, at equal KEGG abundance, may help clarify the across-lineage determinants of protein evolution.

Whether such universal determinants exist remains an open question, although their existence appears likely. This is because a long-standing assumption in the nearly-neutral model of molecular evolution is that species with large effective population sizes (like most microbial organisms) are under strong selection [16–19]. Selection is therefore expected to remove deleterious protein variants more efficiently from prokaryotic populations [20]. Thus purifying selection may have common causes across taxa [18, 21]. The hypothesis that temperature and other environmental variables partly govern the evolutionary constraint of microbial proteins is supported by a varied body of evidence [22–29]. For example, convergent mutations arising in independent *E. coli* populations grown under thermal stress have been observed [30].

Environmental variables may influence microbial protein evolution distinctively. For example, pH may impose different constraints because it is buffered differently among subcellular compartments, but temperature is not [31–35]. Thus, any single-celled organism must encode all its proteins to perform within a single temperature range, whereas different parts of its genome will encode proteins adapted to different pH optima [36].

In this regard, a simple but intriguing question is yet to be addressed: how do environmental temperature and pH increases influence the total number of within-KO sequence variants? Here we have the opportunity to ask this to a metagenomic collection from El Tatio geothermal field, at over 4,200 meters above sea level in the Atacama Desert, Chile. It comprises 17 metagenomes from 13 different thermal locations, with temperature ranging from 45° to 62°C, and pH ranging from 7.2 to 9.3 (Fig. 1). El Tatio geographic isolation, combined with extreme metastable conditions make it a natural laboratory for investigating microbial evolutionary processes [37, 38].

**Figure 1:**
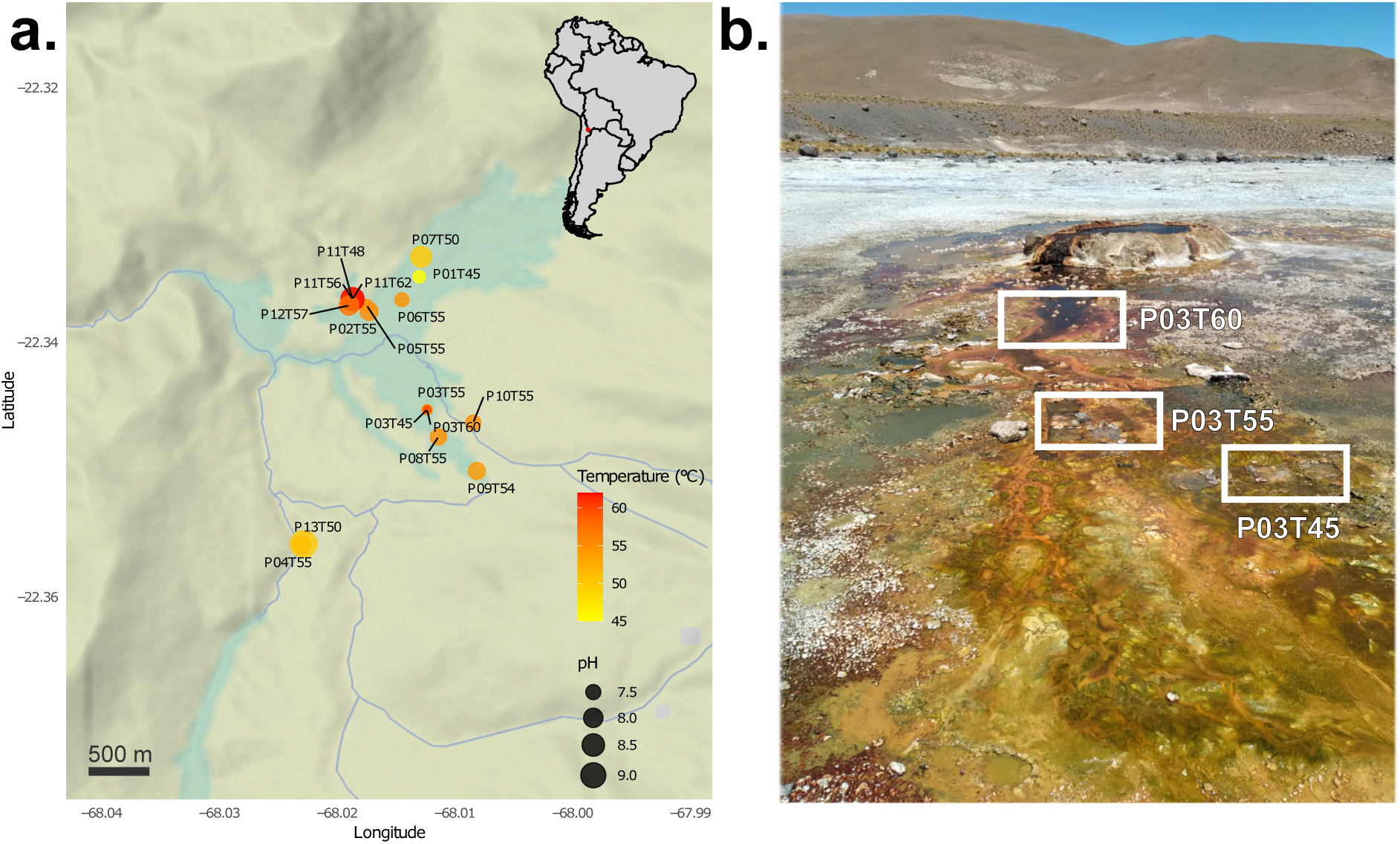
(a). Sampling spots in El Tatio geothermal system. (b). Example of site (P03) in which three different temperatures were sampled (in this case, 45°C, 55°C and 60°C).

Importantly, if temperature or pH universally influence the evolution of these protein repertoires, this signal should emerge clearly in spite of taxonomic heterogeneity among communities. Because our dataset is a community snapshot and cannot trace taxonomic turnover through time, we refrain from inferring direct population genetic parameters. Instead we test whether overall ortholog diversity departs from abundance expectations along temperature/pH gradients.

Ecological dynamics may obviously affect this signal too, but not necessarily spuriously [39]. By favouring taxa carrying particular variants, ecology at the species/strain level can bias which KO sequence variants persist, while HGT allows their fitness effects to be partly decoupled from lineage identity and exposed to selection [40, 41]. Thus, metagenomic sequence diversity may retain a genuine evolutionary signal of temperature/pH-dependent constraints acting at the variant level.

Our results show that, at a given abundance and taxonomic diversity, higher pH was associated with fewer unique sequence contributors per KEGG identifier, whereas higher temperature was associated with more. This effect is found regardless taxonomic trends, although it appears weak. We found partial support for the compartment hypothesis: the effect of pH on residual diversity was distinct in the periplasm. However, we also detected significant heterogeneity between cytoplasmic and membrane proteins in their response to temperature. We therefore conclude that further work is required to falsify the broader compartment hypothesis.

### Within-KO sequence diversity prediction is influenced by temperature categories

In this section, we study the relationship between KEGG abundance and within-KO sequence diversity (hereafter sequence diversity) in our samples (Fig. S1). Sequence diversity is just the count of unique ORFs contributing to a KO ID abundance, where identical predicted proteins are collapsed into a single contributor.

To facilitate the interpretation of results, we collapsed ORFs from different samples corresponding to three temperature bins with equivalent taxonomic diversity (see Fig. S2, Methods).

We checked whether the distributions of sequence diversity (average number of sequences contributing to a KEGG) were different (Fig. S2.1a). All pairwise comparisons were significant (*p* < 0.0001; 45-50°C: median = 9.0, mean = 30.35; 54-55°C: median = 13.0, mean = 38.82; 56-62°C: median = 7.5, mean = 25.41). Thus, the lowest diversity was found in the 56-62°C range.

To better capture the relationship between abundance and sequence diversity, we perfomed bivariate regressions by temperature category. All three linear regressions were significantly different from that of the general dataset (Fig. S2.1b). We checked if the residual distributions on the overall abundance-diversity regression were different. We found every pairwise t-test to be significant (*p* < 0.0001, Fig. S 2.1c). Residuals averaged -2.75 at the lowest temperature range, indicating overprediction (i.e. higher observed than expected once allowing for abundance), peaked 1.45 at the mid range (indicating underprediction), and remained positive (0.35) in the warmest. Medians showed the same trends.

### Sequence diversity prediction is influenced by pH categories

As pH is not correlated with temperature (Fig. S3), we performed the same analysis for three distinct pH categories (Fig. 2.2). The distributions of sequence diversity were likewise different (*p* < 0.0001; 7.2–7.5: median = 7.0, mean = 19.88; 7.6–8.0: median = 10.0, mean = 31.32; 8.1-9.3: median = 13.0, mean = 40.96) (Fig. 2.2a). Without controlling for abundance, the highest diversity is found within the more alkaline pH category. Every regression slope did depart significantly from the general dataset (Fig. 2.2b). While checking the residual distributions, we found that every pairwise t-test was significant (*p* < 0.0001; Fig. 2.2c). The intermediate pH category showed the highest residuals (mean of 1.74). Thus, pH categories also had a genuine effect on sequence diversity when controlling for abundance, although its optimum was reached at the intermediate category (the slope also hints this in Fig. 2.2b, being lower afterwards).

**Figure 2:**
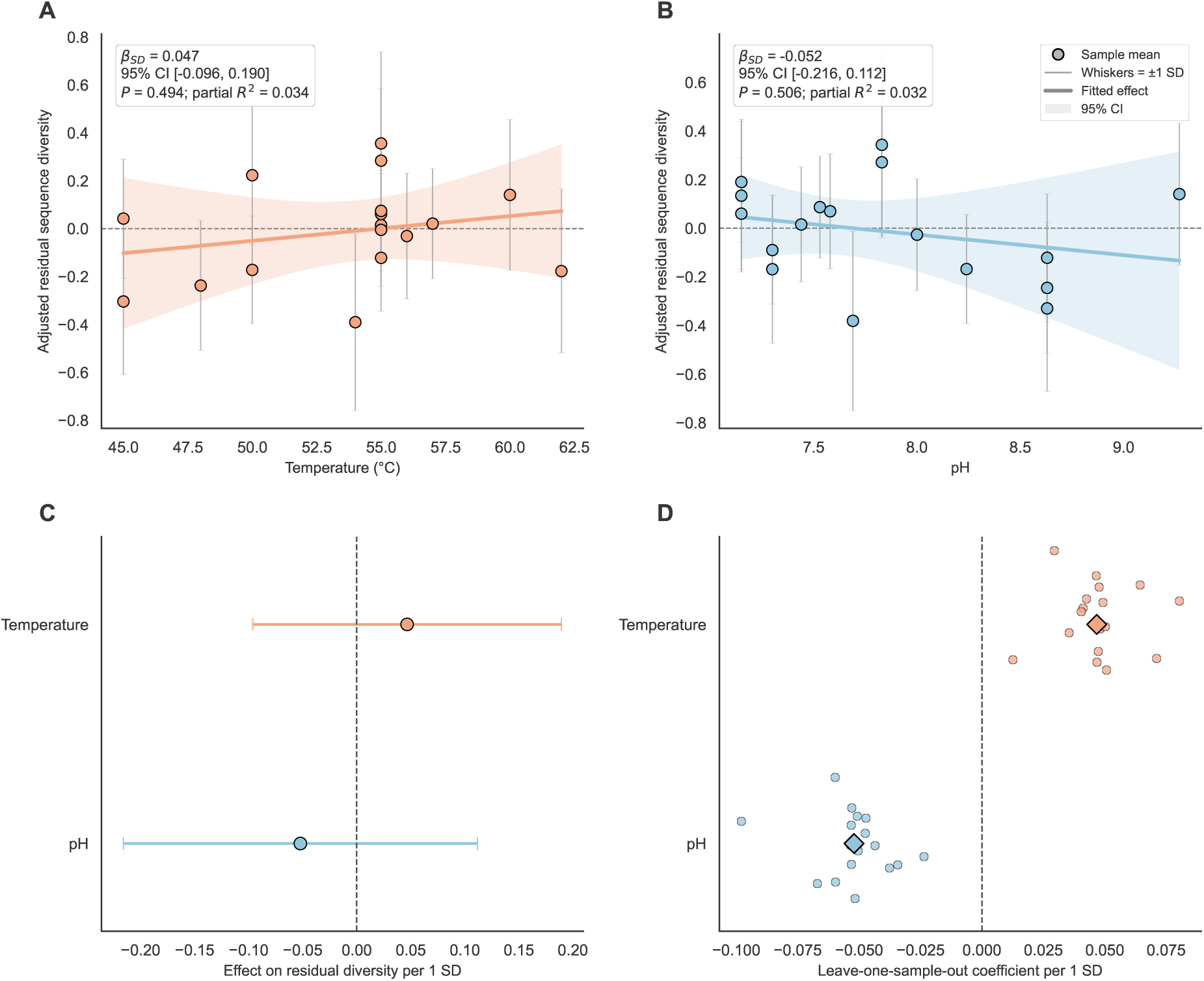
Overall effects of temperature and pH on abundance-controlled sequence diversity. Sequence diversity was quantified for each sample-KEGG combination and adjusted for metagenomic abundance using a KEGG fixed-effect model of sequence diversity against abundance. Residuals were then averaged within each of the 17 independent metagenomes. a–b. Partial effects of temperature and pH, respectively, estimated from a sample-level model including both environmental variables. Points represent sample means and whiskers show ±1 standard deviation across KEGG residuals within each sample. Solid lines indicate fitted effects and shaded regions their 95% confidence intervals. c. Standardised temperature and pH coefficients with 95% confidence intervals. d. leave-one-sample-out estimates obtained by refitting the environmental model after sequentially excluding each sample; diamonds indicate estimates from the complete dataset.

### Higher temperatures predict greater, while alkaline pH predicts lower residual diversity

Although illustrative of the relationship between abundance and community-level within-KEGG sequence diversity, the above results have three important limitations for assessing how well environmental variables fare on predicting sequence diversity: (i) heteroskedasticity persisted in the abundance–diversity regressions, both in raw (*ρ* ≈ 0.63–0.65) and log transformed data (*ρ* ≈ 0.32, BP/White *p* ≪ 0.001), rendering inference from bivariate regressions illegitimate; (ii) we collapsed temperature/pH values into discrete and arbitrary categories for clarity; (iii) unexpected biases in the KEGG distributions might exert random effects on the sequence diversity distributions (e.g. by taxonomic shifts), even when accounting for total KEGG abundance. Thus the observed influence of pH/temperature on sequence diversity could still be spurious.

To confirm or reject our interpretation, we used two complementary approaches: (1) a two-stage analysis of overall abundance-controlled sequence diversity, treating temperature and pH as continuous variables; and (2) a mixed-effects modelling approach in which KEGG identity was included as a random effect (see Methods and Supplementary Information for detailed robustness checks).

Regarding (1), sequence diversity and summed TPM were first calculated for each KEGG identifier within each metagenomic sample. We then fitted a KEGG fixed-effects model relating sequence diversity to abundance (see Methods). The resulting abundance-controlled residuals were averaged within samples and modelled as a function of standardised pH and temperature. The results support an overall weak effect of both variables on prokaryote sequence diversity: higher pH were associated with lower sequence diversity, whereas warmer temperatures with higher (Fig. 2a–c), explaining around 3% of the residual KO variant diversity. This result, although not significant, is robust to sequential exclusion of individual metagenomic samples (Fig. 2d).

Regarding (2), formal inference is based on KEGG-aware mixed models (i.e. KEGG is considered a random effect). Results were consistent as temperature increased sequence diversity (*β* = 0.052, 95% CI: 0.048 to 0.055) and pH reduced it (*β* = −0.050, 95% CI: -0.054 to −0.046) after accounting for abundance and KEGG-level non-independence. A negative-binomial GEE recovered a robust result. Thus, the opposite effects of pH and temperature are not explained by potential KEGG-level skews.

### Temperature/pH – sequence diversity prediction is dependent on subcellular location

We found robust evidence that pH and temperature predict sequence diversity in opposite directions, although their effects appear weak. One explanation is that, in the ranges examined, higher temperature may drive protein diversification (e.g. by fostering neutral mutation rates), whereas higher pH may impose severe functional limitations reducing the set of viable protein solutions.

However, such hypothesis is not testable with our current dataset. One alternative hypothesis is whether the differential control exerted by cells among compartments over some environmental variables, and not others, produces different degrees of protein constraint. This hypothesis might partly explain the opposite departures of pH and temperature. It predicts: (i) compartment-specific *dN*/*dS* after controlling for gene expression (a result already shown in [42]), (ii) compartment-specific sequence diversities, independent of taxonomic preferences, and (iii) that such differences should mainly emerge for variables differentially buffered among compartments, such as pH, viscosity, redox potential, conductivity, etc.

To check whether our data are consistent with predictions (ii) and (iii), we downloaded information of subcellular location from UniProt using the pertinent KEGG identifiers. We next tested whether the abundance-controlled response of sequence diversity to pH and temperature differed among subcellular compartments. Rather than calculating residual diversity in a separate step, we fitted a single interaction model, which estimates how each compartment is influencing the relationship between these two environmental variables and sequence diversity at the same abundance (see Methods).

The results (Fig. 3) indicate that: (1) abundance-controlled sequence diversity is characteristic of each compartment; (2) periplasmic variants become significantly more diverse than variants in other compartments only in relation to pH increases; and (3), unexpectedly, increasing temperature produces greater heterogeneity among compartment-specific coefficients than increasing pH. Thus, temperature and pH show distinct compartment-level associations despite being measured in the same communities.

**Figure 3:**
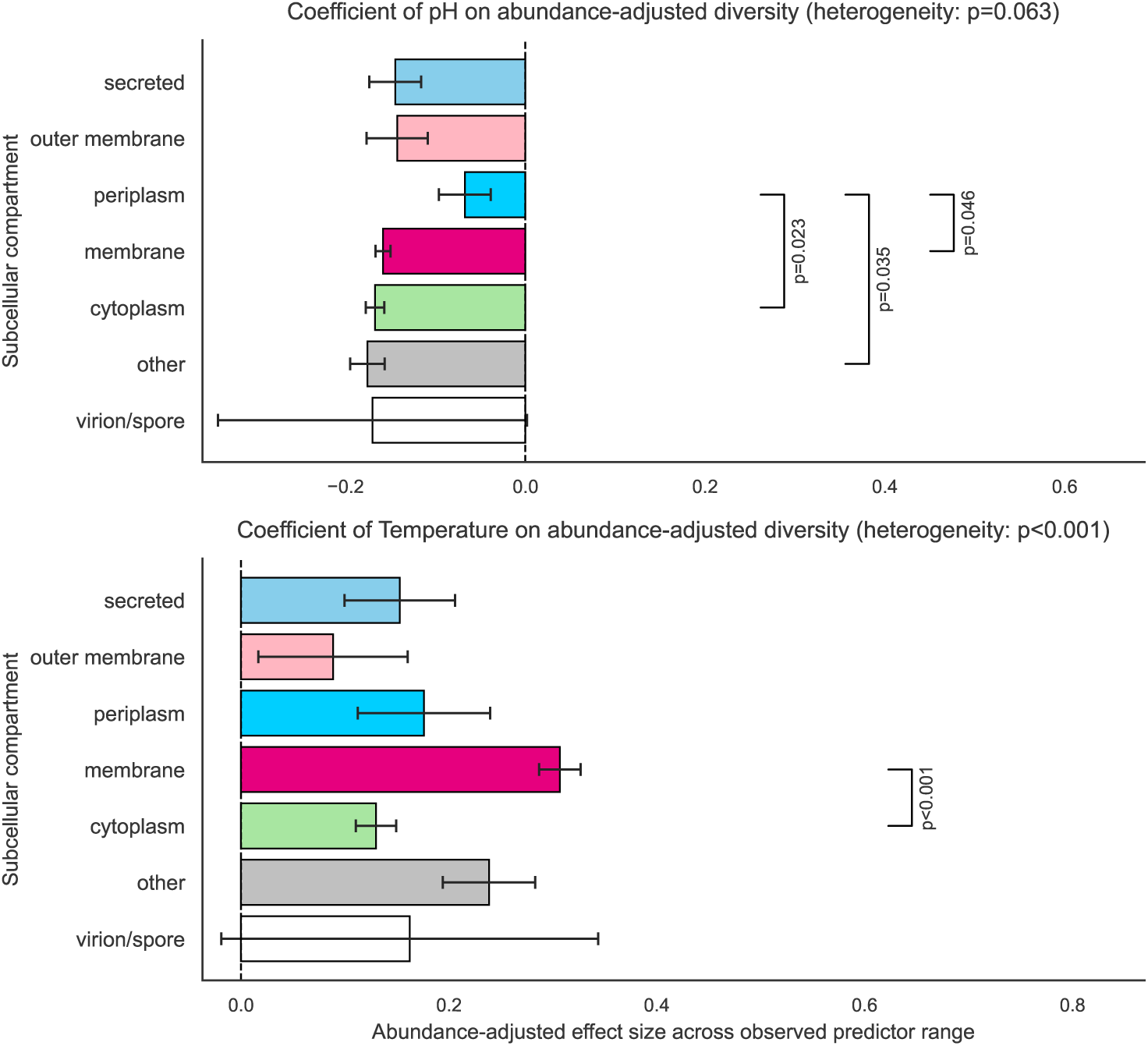
Abundance-adjusted effects of pH and temperature on sequence diversity across subcellular compartments. Bars show compartment-specific coefficients for pH (above panel) and temperature (below panel), estimated from a single interaction model, at equal abundance. Predictors were scaled across their observed ranges, such that each coefficient represents the expected change in sequence diversity from the minimum to the maximum observed value, at fixed abundance. Global Wald tests assessed heterogeneity among compartment-specific slopes. Brackets indicate significant pairwise differences between slopes, with *P* values adjusted by the Holm method. Standard errors were clustered by KEGG identifier.

These results support that periplasmic sequence diversity is distinctively affected by the external pH oscillations across taxa, and that the universally negative influence of pH on residual diversity appears to be constant in all the other compartments. We nonetheless find significant heterogeneity in residual diversity between cytoplasm and membrane with temperature increases (3). At first sight, this point is contradictory with prediction (iii), which makes us reject a naive compartment hypothesis. This point will be revisited in more detail in the discussion.

### Most abundant KEGGs are depleted in warmer environments, but overrepresented in alkaline pH samples

We found strong evidence that higher temperature is associated with greater abundance-controlled sequence diversity, and alkaline pH with diminished diversity (in the ranges examined). This relationship appears to be dependent on compartment, too. But, why is this so? We expected a reduction in sequence diversity with temperature (see Discussion), but we observe the opposite.

We wondered whether this apparent contradiction could be explained by a preferential depletion of highly abundant KEGG functions in warmer communities. Under this scenario, warmer environments would contain fewer highly represented functions, reducing total sequence diversity, while the KEGG functions that remain would contain more sequence variants than expected from their abundance (e.g. by enhancing mutation). To test this we have analysed both halves of the abundance distribution: the top 50% and the bottom 50% abundant KEGGs (Fig. 4).

**Figure 4:**
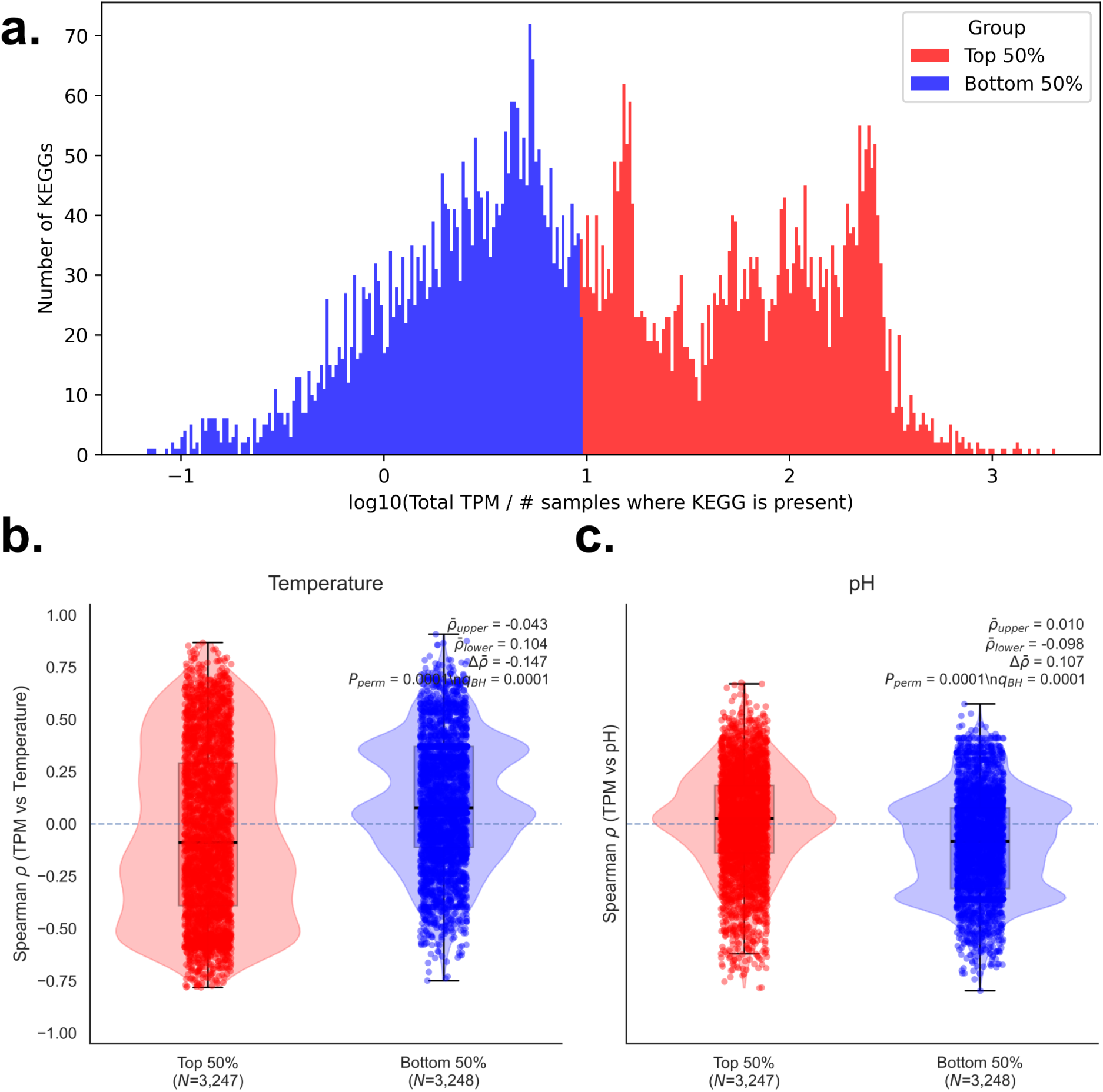
(a). Distribution of KEGG abundance. To make a conservative test, we split the distribution in two, to search if there is a characteristic behavior of the most abundant fraction with the variables. (b). TPM was summed for each metagenome per KEGG combination, with undetected combinations assigned TPM =0. KEGGs were divided into the top (red) and bottom (blue) abundance halves across all metagenomes. Each point is the Spearman correlation between one KEGG’s abundance and temperature (b) or pH (c). Violins and boxplots show the coefficient distributions. P-values derive from permutation tests of the difference in mean *ρ* between abundance classes; q-values were Benjamini–Hochberg adjusted. These results are supported by 10% top comparison (Fig. S7) and regressions of KEGG loses, in Figures S5 and S6.

In regard to temperature, we found that the more abundant KEGGs were significantly depleted in warmer samples (*p* = 0.0001, -0.043 Spearman’s *ρ*), (Fig. 4b). We also performed a Monte Carlo test (10,000 extractions) to visualize how enriched in negative correlations with temperature are the more abundant KEGGs (observed mean *ρ* (Top 50%): -0.0814; mean of random draws: -0.0645 ± 0.0034; empirical p-value (2-sided): 0.0002; Fig. S5a).

But are the most abundant KEGGs disproportionately absent from warmer metagenomes? To test this, we regressed each KEGG’s total abundance across the dataset against the mean temperature of the metagenomes in which that KEGG was undetected after summing TPM across all taxa. We found that the slope was positive and significant (slope = 0.45; p = 0.0125; Fig. S5b). As total abundance of a KEGG increases, it makes more probable for that KEGG to be lost at higher temperatures.

These results indicate that, in absolute values, sequence diversity is of lesser magnitude in hotter springs (Fig. S2.1a), but this is only caused by the fact that low-abundance KEGGs are preferred (Figs. 4a, S5) – corresponding high diversities are not achieved, as anticipated in Figure S2.1b.

When interrogating the data for increasing pH values, we find the opposite behavior: the more abundant functions are more abundant in alkaline springs (*p* < 0.0001; 0.01 Spearman’s *ρ* Fig. 4b). Again, Monte Carlo tests are in agreement with the latter (observed mean *ρ* (Top 50%): 0.0217; mean of random draws: 0.0072 ± 0.0034; empirical p-value (2-sided): <0.000; Fig. S6a). We performed the regression to ascertain in which pH a KEGG is lost as its total abundance increases. This time we found that the slope was slightly negative (slope = -0.096; p = 0.02; Fig. S6b). As abundance increases, the KEGG loss probability for higher pH values is decreasing. Surprisingly, this represents an opposite response to that induced by temperature. This is especially meaningful given that pH is not correlated with temperature in our dataset (Fig. S3).

Our results are robust when comparing the more abundant KEGG fractions (top 10%). We confirmed that they were significantly different (mean *ρ* (Temperature): -0.08 ± 0.15; mean *ρ* (pH): 0.02 ± 0.14; t = –28.14, p = 0.000; Fig. S7).

### Proteins are shorter in warmer and alkaline environments

A complementary test is checking whether discrepancies in protein length with increasing temperature and pH match patterns of diversity.

Protein length distribution is considered mostly uniform across the tree of life [43]. As our metagenomes include proteins from astranged branches of the tree of life and of equivalent taxonomic richness (Fig. S8), our protein length distributions should remain fairly uniform across pH and temperature categories, wherever they do not have a true influence on length.

Contig characteristics were comparable among environmental categories, with no detectable differences in contig length, N50 or coverage across temperature (P=0.650–0.994) or pH bins (P=0.299–0.822).

When interrogating our data, the results showed clear trends (Fig. 5). Significantly shorter protein lengths corresponded only to the warmest temperature category (average aa: 45-50°C = 164.61; 54-55°C = 164.79; 56-62°C = 148.66; pairwise t-test p-values indicated in Fig. 5a). Likewise, the fraction of Long Proteins (LP; > 1, 000 aa) was significantly depleted as temperature increased (Fig. 5b). These results are in agreement with previous observations in phylogenetically reduced groups [44, 45].

**Figure 5:**
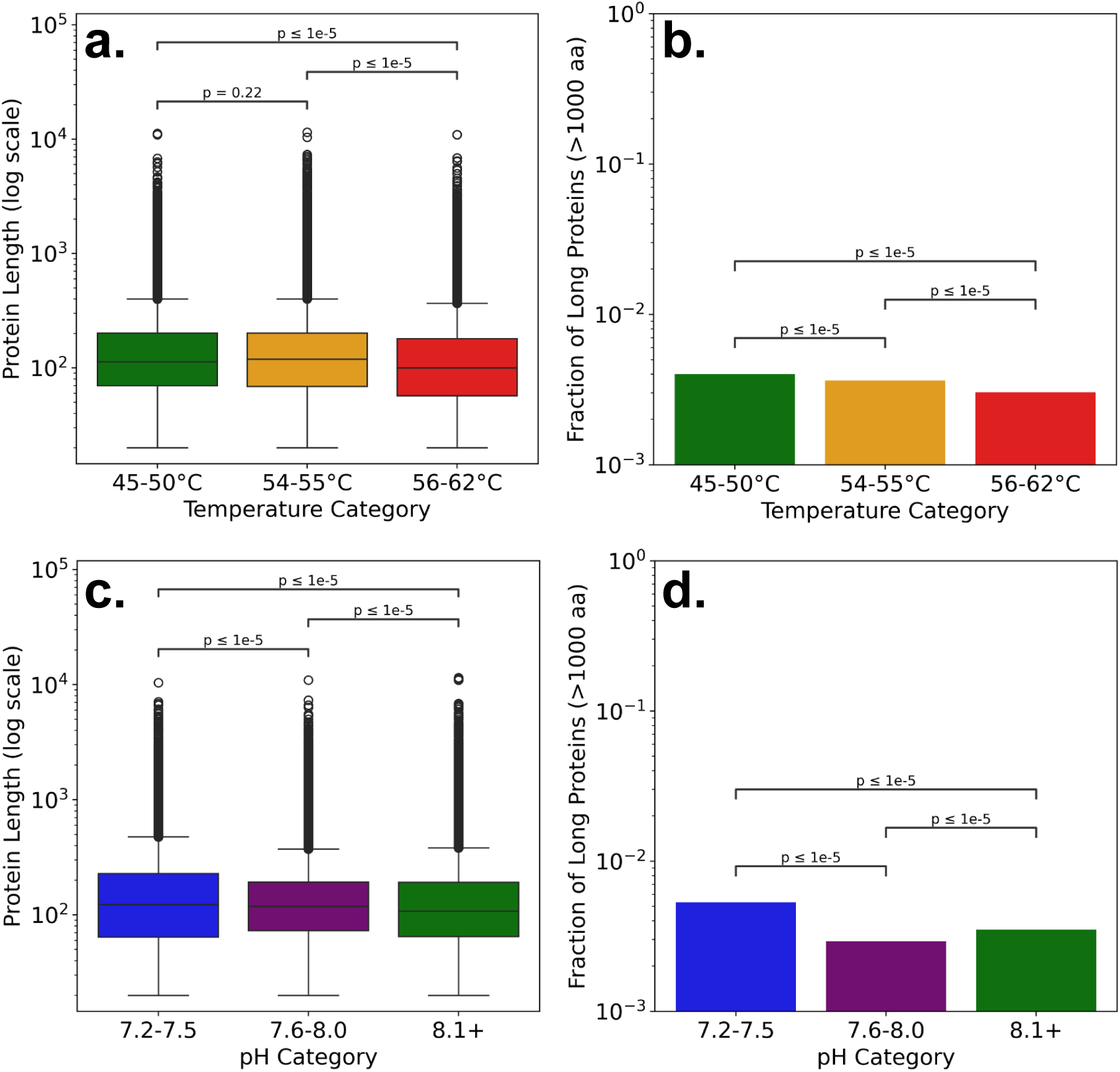
a. Distributions of protein length by temperature categories (all domains). Only higher temperature bin (red) was shown significantly shorter proteins (paired t-tests). b. Log fraction of proteins spanning more than 1,000 aa classified by temperature. The distributions showed a significant depletion of long proteins that correspond to higher temperatures (confidence intervals are negligible). c. Distributions of protein length by pH categories (all domains). In this case, every category showed significant differences. d. Log fraction of proteins spanning more than 1,000 aa classified by pH category. This result suggest that enrichment of long proteins is anticipated in metagenomes retrieved from an environment outside the 7.6–8.0 pH range (purple), and that this also applies to more alkaline environments.

Interestingly, while a more alkaline pH is associated with a shorter protein length (average aa: 7.2-7.5 = 177.91; 7.6-8.0 = 160.37 8.1-9.3 = 157.16; Fig. 5c), LP enrichment was found in both the more neutral and alkaline ranges.

## Discussion

Here we explored how the general relationship of sequence diversity and metagenomic abundance may be broadly influenced by temperature and pH in a metagenomic collection retrieved from hot spring communities. We here have shown complementary results supporting the interpretation that pH and temperature are weak, but probably legitimate predictors of protein evolution – although with contrary sign. Within the range of values inspected, increasing pH values diminish, while increasing temperature foster the KO sequence diversity over metastable populations. These effects are found when interrogating the residual diversities of unique sequence contributors not explained by KEGG abundance.

Importantly, the observation of higher residual diversity is not explained by characteristic taxonomic richness in different temperature or pH arbitrary categories, as no significant differences were found among these (Fig. S8).

### Is adaptation to higher temperatures driving genome size reduction?

On the one hand, the warmer temperatures in our dataset underpredicted the number of sequences found. Thus we consistently observed higher variant diversity, ensuring equal KEGG abundance, in metagenomes from warmer springs. We had expected the opposite pattern, as higher temperatures limit taxonomic diversity and should similarly narrow the range of sequence variants compatible with protein function.

On the other hand, this result appear to contradict the observation that thermophilic organisms are prone to genome size reduction through gene loss [46–49]. Genome size reduction in metabolisms adapted to higher temperatures is independently predicted by the biosynthetic model, in which higher body temperatures cause adaptive reductions in genome size [50].

However, our results do not argue against reduced genome size as a consequence of thermal adaptation. We can reconcile our results with previous observations by simply assuming that genome reduction is achieved non-randomly. Thus genome reduction may result from the preferential loss of variants belonging to more dispensable, but highly represented KEGG functions; from temperature-associated shortening of coding sequences; or from some combination of both. Likewise, any contribution of sequence shortening (Fig. 5) to genome size reduction would be expected to concentrate disproportionately on dispensable regions [51].

One hypothesis is that warmer environments reduce the total net number (because of abundant KEGG depletion) and/or length of gene variants while simultaneously increasing the diversity of variants represented across taxa within the KEGG functions retained in the community. Indeed, genome reduction has been associated with accelerated protein evolution in free-living prokaryotes, showing that genome streamlining need not entail reduced sequence variation [52]. This possibility is also consistent with observations in particular *Fischerella thermalis* ecotypes where divergence among orthologs increased with temperature and exclusive genes appeared at 66 ^◦^C [53]. This remains nonetheless speculative.

### The “confined evolution” hypothesis

We further tested whether abundance-controlled KO sequence diversity differs among subcellular compartments, and whether its association with pH or temperature depends on protein localisation. Microorganisms cannot maintain different temperatures across their subcellular compartments, whereas environmental pH is buffered to different degrees across them. We therefore expected compartment-specific responses to be more pronounced for pH than for temperature.

Our results only partly support this expectation. The association between residual sequence diversity and temperature varied more strongly among compartments overall (Fig. 3). Pairwise comparisons showed that cytoplasmic and membrane proteins strongly differed in their response to temperature. By contrast, periplasmic sequences differed significantly from other compartments only in their response to pH, consistent with the more direct exposure of the periplasm to extracellular proton concentrations [36].

Thus, there is a discrepancy between the model predictions and the observed features. One simple explanation imply that the model is naive. For example, we assume that temperature does not differ among subcellular locations because it is not actively regulated by the cell. An alternative model could incorporate cell size and geometry as relevant factors, as differences in these properties among species may modulate transient exposure to environmental temperature fluctuations. However, sustained intracellular thermal gradients are unlikely at the scale of prokaryote cells.

Another plausible explanation is that the same environmental temperature imposes different molecular constraints across compartments. We speculate that the observed discrepancies between cytoplasmic coefficients may reflect differences in exposure to thermal shock, structural constraints, or the distinct effects of temperature on membrane-associated and intracellular metabolic processes.

While our results provide preliminary support for the intuition that compartment-dependent buffering may contribute to differential responses to uncontrolled environmental fluctuations, further work is required to test the broader implications of such a model.

For example, if the confinement hypothesis were correct, evolutionary rates should also differ among compartments across species, provided that equivalent compartments exert comparable control over the same environmental variables. Evidence consistent with this prediction is provided for *E. coli* and *S. cerevisiae* in [42]. Contrary to the extended complexity hypothesis [54], differences in evolutionary rates or sequence diversities among subcellular localisations would not primarily reflect the number of functional interactions of a protein, but rather gene expression, essentiality, and subcellular location. We content that the effect of location respond to the differential ability of cells to buffer environmental stresses from one compartment to another.

### Limitations

In this exploratory study, we used the number of distinct sequences assigned to each KEGG function as a coarse proxy for protein diversification. This metric does not distinguish among mutation, gene duplication, horizontal transfer or taxonomic turnover. Moreover, KEGG groups represent functional orthology and do not necessarily comprise strict genealogical orthologues; some of the observed diversity may therefore reflect paralogous or evolutionarily heterogeneous sequences. Our analysis should consequently be regarded as a first methodological approximation to within-function sequence diversification.

The effects of pH and temperature cannot be considered fully independent (although noncorrelative; Fig. S3), as each sample reflects both environmental influences at the same time. The individual contributions of pH and temperature therefore cannot be completely disentangled from observational data alone. The number of truly independent samples is also low (n=17). Future experimental work should manipulate these variables independently.

Another consideration would be that the sequencing depth of samples might be different, or that our approach considered only annotated ORFs (avoiding broader measures of sequence diversity). Nevertheless, (i) we used TPM averages per KEGG per condition (reducing detection noise); (ii) the non-pareil analysis exhibited sequencing coverage (C values) above the 95% threshold (range: 96.34–99.52%) for all but one sample (93.21%), indicating that the observed patterns likely reflect the true community composition; (iii) we made robustness checks (see Supplementary Information).

More importantly, because residual within-KO diversity was quantified at the whole-metagenome level, the observed patterns cannot be interpreted independently of taxonomic composition or phylogenetic structure. Taxonomic turnover may contribute additional variance, but should generate a systematic environmental signal only insofar as community composition strongly covaries with temperature or pH. Taxonomic composition showed little covariance with pH (Bray–Curtis *R*^2^ = 0.028, *P* = 0.858; Jaccard *R*^2^ = 0.057, *P* = 0.594) and only modest covariance with temperature (*R*^2^ = 0.191, *P* = 0.009; *R*^2^ = 0.104, *P* = 0.011, respectively), the latter becoming non-significant after exclusion of the warmest bin.

We argue that, although changes in taxonomic composition across environmental conditions are expected to partly explain KO variant patterns, those compositional changes are themselves structured by temperature and pH at the broader community scale.

Finally, our results only encompassed hydrothermal organisms in particular, limited conditions. Because the dataset was dominated by bacterial and viral ORFs (Fig. S1), the conclusions may not extend to other domains.

## Methods

### Sampling & sequencing

Samples were obtained from 13 hot spring manifestations of the El Tatio geothermal field in January 2020, which their metadata and sequences were reported in previous studies [55, 56] and are available in Table S1. Briefly, triplicate samples of approximately 2 ml were collected with a punch from each microbial mat, kept in cryogenic vials containing RNAlater (Thermo Fisher Scientific, Vilnius, Lithuania), and then stored at -80°C until DNA extraction. DNA extractions were performed according an optimized protocol for microbial mat samples from hot springs [57]. An equimolar amount of DNA (400 ng) from each replicate was pooled and sequenced at the Roy J. Carver Biotechnology Center (University of Illinois at Urbana-Champaign, IL, USA with the Illumina NovaSeq 6000 platform (S1 flowcell, 2×150 bp).

### Metagenomic analysis

The resulting fastq files for metagenomes were filtered for quality using Cutadapt [58] with the parameters: paired-end mode, a perfect match of at least 10 bp (-O 10) against the standard Illumina adapters, hard trimming of the first 10 to 15 bases of the 5’ (-u 10/15) and 3’ (-U 10/15) ends depending on quality and base distributions, removal of N-containing reads (–trim-n 1) and retention of only sequences longer than 50 bp (-m 50). De novo assemblies of trimmed reads were generated using SPAdes v3.10.1 [59] with the –meta setup. Sequencing effort was assessed with the Nonpareil v3.401 software [60] with the -T alignment and -f fasta options.

The CDS were predicted with the Prodigal V2.6.3 software [61] with the -p meta option. Therefore, at the wider level, all the contigs were classified with geNomad v1.8.1 (using the end-to-end option; [62]) in viral and plasmid contigs, and then the unclassified >1000 pb (–minsize 1000) contigs were assigned to prokaryotes and eukaryotes with whokaryote software [63], leaving all other contigs as unclassified. Then, the taxonomic affiliation of prokaryotic and plasmid contigs and proteins was obtained with the refineM tool [64], using the entire assemblies as input for the modules scaffold stats and the assignment with the module taxon profile using also the previously predicted CDS and a custom protein/taxonomic diamond v2.1.9.163 [65] database with all the proteins from the representative genomes from the GTDB R220 (available at https://data.ace.uq.edu.au/public/gtdb/data/releases/release220/220.0/genomic_files_reps/) and formatted as required with a custom python script.

To functionally annotate the predicted ORFs, DRAM v1.5.0 software [66] was used with the annotate genes and distill options and retrieving the results from the KOfam [67] database. Then, reads were aligned to the nucleotide sequences of the predicted ORFs with the coverM v0.6.1 (https://github.com/wwood/CoverM; [68]) with the following parameters: coverm contig -p bwa-mem –methods covered bases mean rpkm tpm length count covered bases –min-read-percent-identity 95 –min-read-aligned-percent 50 and –min-covered-fraction 0 against their respective metagenomes. The functional, taxonomic, protein length, and abundance information was parsed to each contig and each ORF with custom bash and python scripts.

### Modelling of sequence diversity

For each sample we computed the total transcript abundance (TPM) and the number of open reading frames (ORFs). In total, the dataset comprised 17 samples with a cumulative TPM of 17,000,193 and 1,381,486 ORFs.

To assess how environmental variables influence sequence diversity beyond that expected from metagenomic abundance, we implemented a two-stage linear model. This approach separates the influence of abundance on sequence diversity from that of the two environmental variables tested, allowing us to estimate whether pH and temperature are associated with consistent deviations in the number of distinct sequences assigned to each KEGG identifier.

In the first stage, observations were aggregated by sample and KEGG identifier. Sequence diversity was defined as the number of distinct sequence clusters associated with each KEGG identifier within each sample, whereas abundance was calculated as the summed TPM associated with that KO within the same sample. We then fitted the following KEGG fixed-effects model:

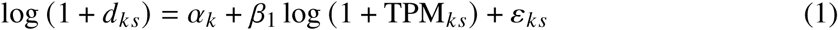

where *d*_*ks*_ is the sequence diversity of KEGG identifier *k* in sample *s*, TPM_*ks*_ is its summed abundance, and *α*_*k*_ is a KEGG-specific intercept accounting for intrinsic differences in sequence diversity among gene functions. The model was fitted by within-KEGG transformation, which is algebraically equivalent to including a fixed intercept for each KEGG identifier.

The residuals from this model represent variation in by-KEGG sequence diversity not explained by abundance. Positive residuals indicate greater sequence retrieval than expected from abundance, whereas negative residuals indicate underrepresentation. Residuals were subsequently averaged across KEGG identifiers within each sample, yielding one estimate of abundance-controlled sequence diversity per independent metagenomic sample.

In the second stage, we modelled mean residual diversity as a function of environmental conditions:

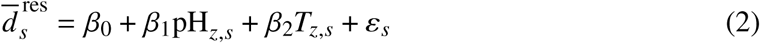

where 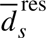 is the mean residual diversity of sample *s*, and pH_*z,s*_ and *T_z,s_* are the standardised pH and temperature values of that sample. Both environmental variables were treated as continuous predictors and standardised to a mean of zero and a standard deviation of one. Their coefficients therefore represent the average change in residual sequence diversity associated with a one-standard-deviation increase in each environmental variable, conditional on the other. This approach represents a specific application of the methodology developed in [69]. Models were fitted by ordinary least squares using the “statsmodels” library in Python.

### Temperature, pH, and subcellular categorization

As described, TPM values were normalized by the number of unique samples within each environmental category (temperature or pH). Environmental variables were discretized into arbitrary intervals: 7.2–7.5, 7.6–8.0, and 8.1–9.3 for pH; and 45–50^◦^C, 54–55^◦^C, and 56–62^◦^C for temperature. For display, ranges were rounded to the nearest tenth (e.g. 9.3 corresponds to 9.27). These bins were selected to reflect with clarity the trends in sequence diversity variation while ensuring an approximately even distribution of metagenomes across categories. Subcellular localization was assigned by querying UniProt with cleaned protein descriptions from KEGG identifiers. For each entry, the first UniProt match was retrieved and its annotated subcellular compartment extracted from the JSON API. We collapsed localisations for clarity and statistical resolution. “Membrane” corresponds to any membrane that is not “outer”. We removed ambiguous fetches.

Further methods are available in the Supplementary Information.

## Data availability

The custom code to generate the figures and analyses (with specific methods) are available at 10.5281/zenodo.15701078. The metagenomic raw reads can be found under the NCBI BioProject accession PRJNA858297 (https://www.ncbi.nlm.nih.gov/bioproject/PRJNA858297/).

## Acknowledgements

We want to thank Alvar Lavin, Pablo Yubero, Juan F. Poyatos, Sofia Radrizzani, and Laurence D. Hurst for a variety of inputs helpful to this work. We thank the organizing committee and members of both the National Network of Extremophiles (RedEx) and 2025 Microbiology Society Athlone Annual Meeting for helpful discussions.

## Authors’ Contributions

J.R.S.: Conceptualization, Methodology, Software, Formal Analysis, Investigation, Validation, Resources, Data Curation, Visualization, Writing - Original Draft, Writing – Review & Editing, Supervision; J.A.: Methodology, Validation, Resources, Data Curation, Writing – Review & Editing, Visualization; B.D.: Validation, Resources, Writing – Review & Editing; J.T.: Resources, Project administration, Funding acquisition, Writing – Review & Editing; C.P.A.: Conceptualization, Investigation, Validation, Writing – Review & Editing, Supervision, Project administration, Funding acquisition.

