## Supplementary Information for "Across-lineage prediction of within-KO sequence diversity by temperature and pH differs among subcellular compartments in hydrothermal spring communities"

### Temperature and pH universally govern protein diversity in hydrothermal spring communities, but they do so differently

Rivas-Santisteban *et al.* (2025)

---

**Methods: Assessment of KEGG depletion**

To explore how KEGG functions are depleted across different environmental conditions, we analyzed TPM-normalized abundance data for genes annotated with KEGG identifiers. We ranked all KEGGs by their total TPM across the dataset and divided them into two groups: the top 50% (high-abundance) and the bottom 50% (low-abundance). Within each group, we calculated Spearman correlations between environmental variables (like temperature and pH) and the normalized TPM values for each KEGG. We then compared the distributions of these correlations using Welch's *t*-test. To visualize the results, we used violin plots shown in Figure 4, interpreting the differences in correlation strength as an indication of whether abundant and rare KEGGs respond differently to environmental changes.

**Methods: Protein length distributions**

To examine how protein length varies across environmental conditions, we analyzed amino acid sequence files (.faa) for each metagenome. These files were named according to their sample context and temperature, which allowed us to match them with relevant metadata. We extracted each protein sequence line by line and recorded its length in amino acids. Temperature values were pulled from the filenames using regular expressions and assigned to predefined categories. pH values were retrieved separately from a metadata table and matched to each sample using unique identifiers. For each protein, we compiled a table that included its length, associated temperature and pH values (and their categories), context ID, and the original filename.

**Methods: Abundance-adjusted environmental coefficients among compartments**

We tested whether the relationship between environmental variation and sequence diversity differed among the established subcellular compartments. KEGG observations were first aggregated by KEGG identifier, compartment, pH, and temperature. Sequence diversity was estimated as before.

We fitted a single abundance-adjusted interaction model:

$$\log(1 + D) \sim C + C \times [\log(1 + A) + pH + T]$$

where D is sequence diversity, A is total abundance, C is compartment, and T is temperature. pH and temperature were scaled from 0 to 1 across their observed ranges, so their coefficients represent the expected change in abundance-adjusted sequence diversity across the full observed environmental range. The inclusion of compartment-by-environment interactions allowed each compartment to have its own pH and temperature coefficient.

We tested overall heterogeneity among compartments using Wald tests on the compartment-by-environment interaction terms. Pairwise differences between compartment-specific environmental coefficients were evaluated as linear contrasts from the same global model, with Holm correction for multiple testing. Standard errors were clustered by KEGG identifier to account for repeated observations of the same functional category across environments and compartments.

**Methods: robustness checks**

Robustness was further assessed using three complementary analyses. First, we performed a leave-one-sample-out analysis by sequentially excluding each metagenomic sample and refitting the sample-level environmental model.

Second, using all ORFs aggregated by metagenome and KEGG identifier, we fitted the following KEGG-aware mixed-effects model:

$$\log(1 + \text{gene count}) \sim \log(1 + \text{sum TPM}) + \text{pH}_z + \text{Temperature}_z + (1 \mid \text{KEGG})$$

where KEGG identity was included as a random intercept. From a narrative viewpoint, this model implies: After controlling for abundance and baseline differences among KEGG functions, are pH and temperature associated with the observed number of sequences?

Finally, we fitted a negative-binomial generalised estimating equation to the untransformed gene counts, with log(1+ sum TPM), pH, and temperature as predictors. Observations were clustered by KEGG identity using an exchangeable working correlation structure, thereby accounting for repeated measurements of the same KEGG identifier across metagenomes. This analysis tested whether the inferred environmental effects were robust to modelling sequence diversity directly as count data rather than after log transformation.

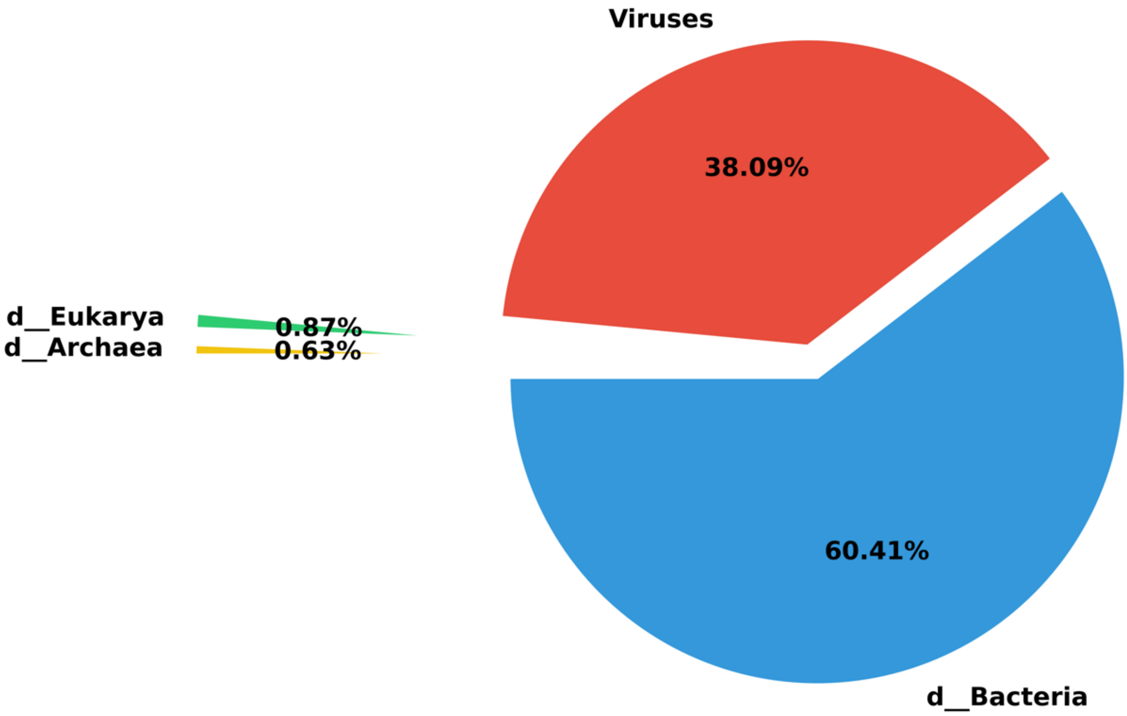


Fig. S1. Taxonomic distribution of mapped ORFs across El Tatio retrieved metagenomes. A total of 6,475 KEGG orthologies were shared across multiple domains. Among domain-specific sets, Bacteria harboured the greatest number of unique KEGGs (6,168), followed by Acytota (2,643), Eukarya (2,582), and Archaea (1,321).

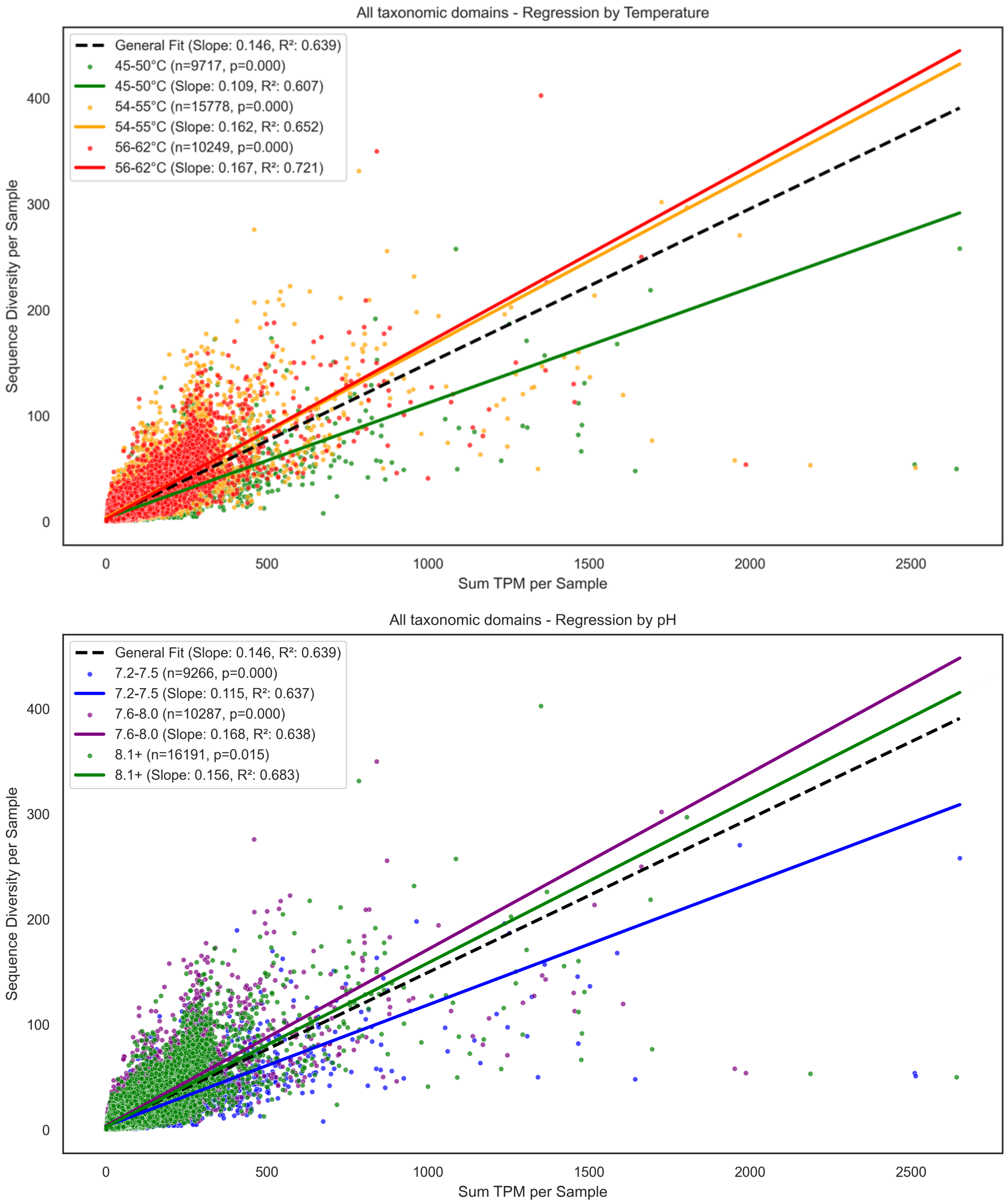


Fig. S2. Relationship between abundance and diversity collapsed by KEGG (all taxa in Fig. S1 including viruses) is dependent on environmental data. Coloured regressions represent different subsets according to arbitrary bins on temperatura and pH. Further details are provided below for exclusive bacterial ORF regressions.

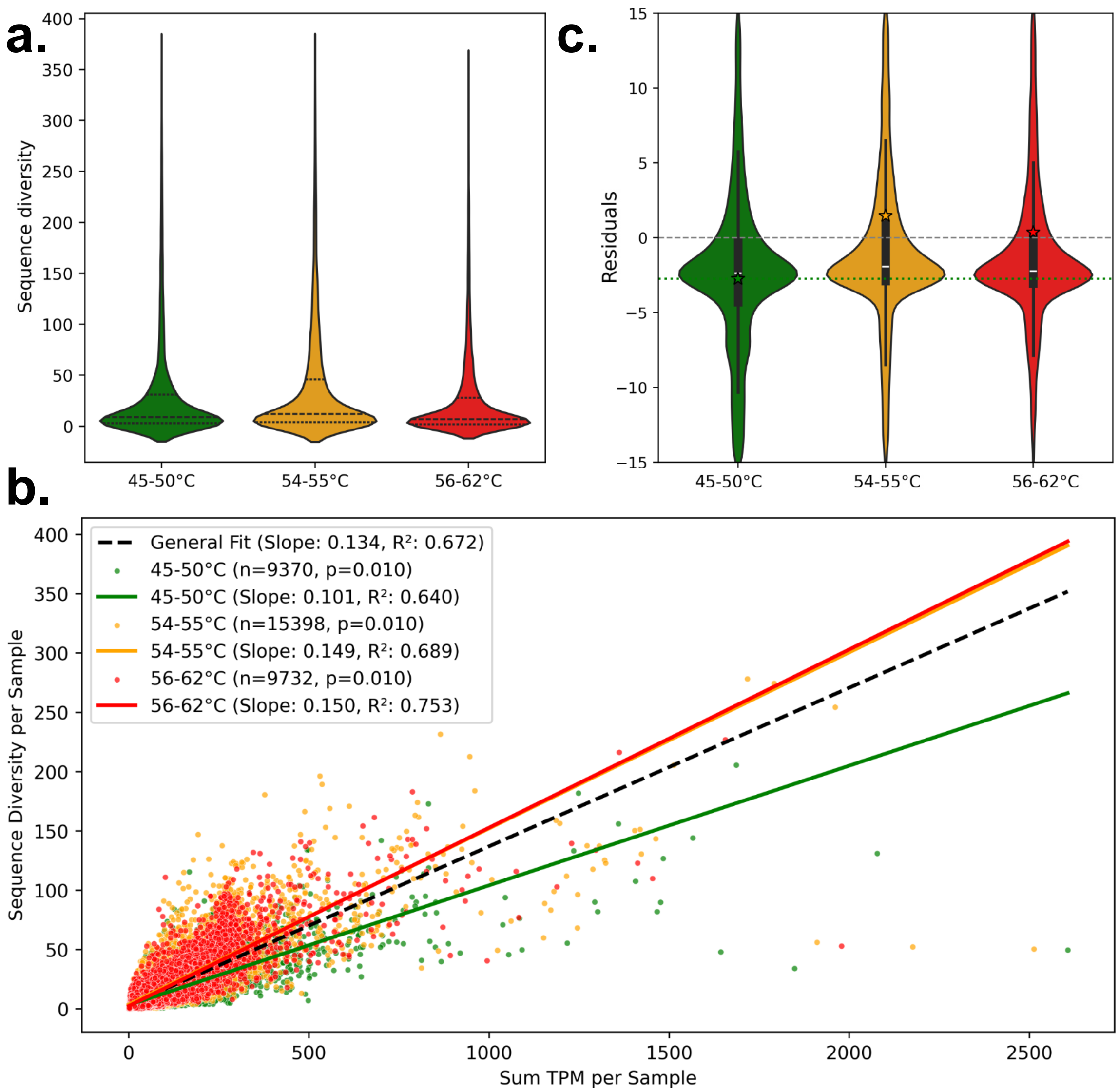


Fig. S2.1. a. Distributions of sequence diversity (predicted bacterial ORFs for the same KEGG) are significantly different in all temperature categories tested (paired t-tests, $p < 0.0001$). This result is nearly identical to the overall abundance-diversity relationship A priori, the lowest diversity of sequences per function is found in the 56-62ºC category. b. General and specific linear regressions among temperature categories. The explanatory power of genomic abundance consistently increases with temperature. Every slope differs from the general regression (10,000 iterations, p-values indicated). We observe that sequence diversity grows faster with abundance, but warmer temperatures are depleted of abundant functions. c. After allowing for abundance, residual distributions are found to be significantly different across temperatures ($p < 0.0001$), with the 54-55ºC category containing the higher by-function diversity. Stars indicate means. Range limited to ±15 for visualization. The variance is gradually reduced with temperature, which explains the better linear fit within warmer metagenomes.

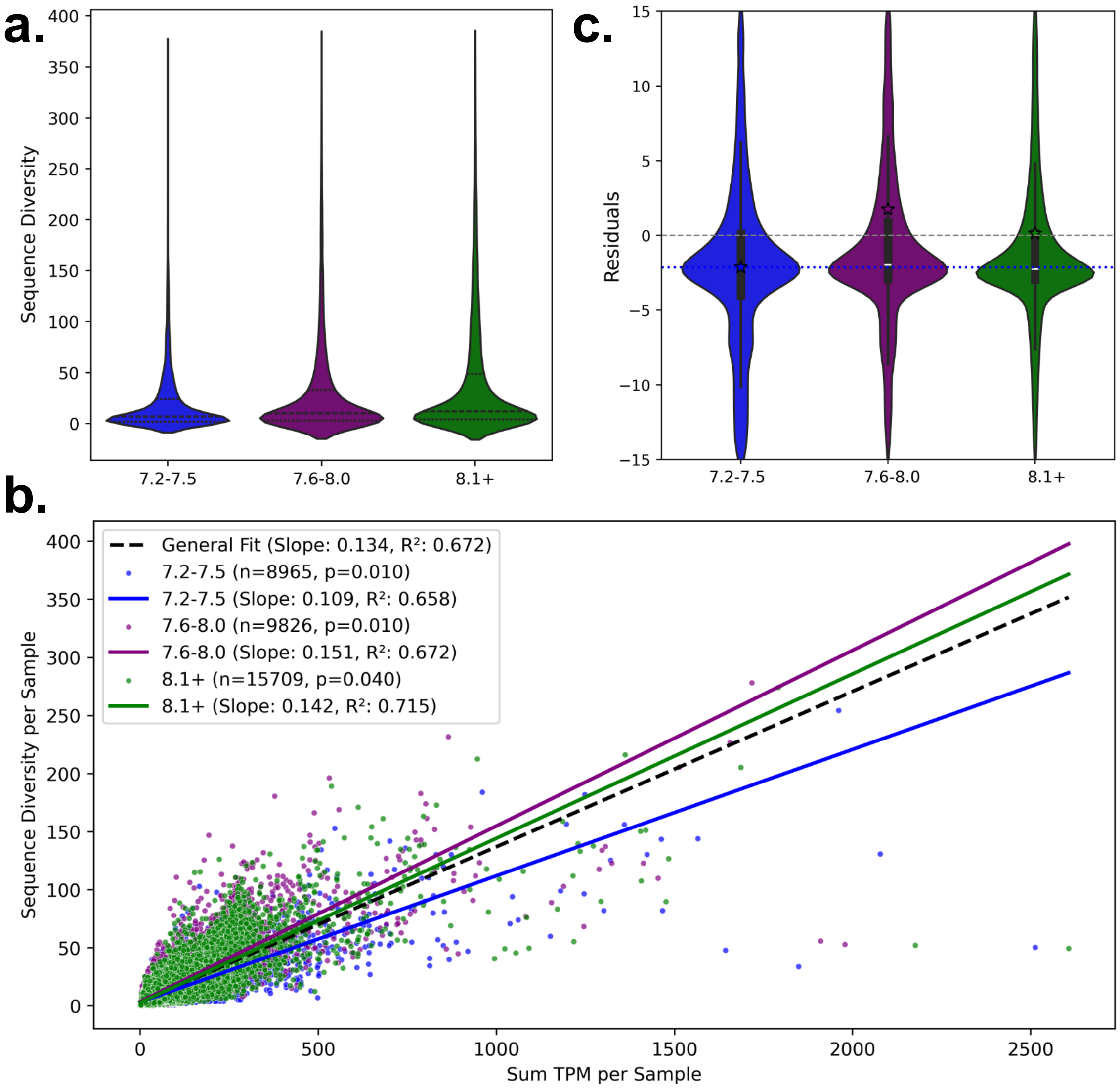


Fig. S2.2. a. Distributions of gene counts in Bacteria are also different among pH categories, and near-identical to the overall abundance-diversity relationship (paired t-tests, $p < 0.0001$). b. General and specific linear regressions among temperature classes. In this case, the explanatory power of genomic abundance is barely increasing with pH. The 8.1-9.3 pH range slope is not differing from the general regression. c. Higher pH residual distribution is likewise significantly lower ($p < 0.0001$), although the higher pH range has a lower significance ($p = 0.04$). Stars indicate means. Range limited to ±15 for visualization. Unlike temperature, residuals show equal variance as pH increases.

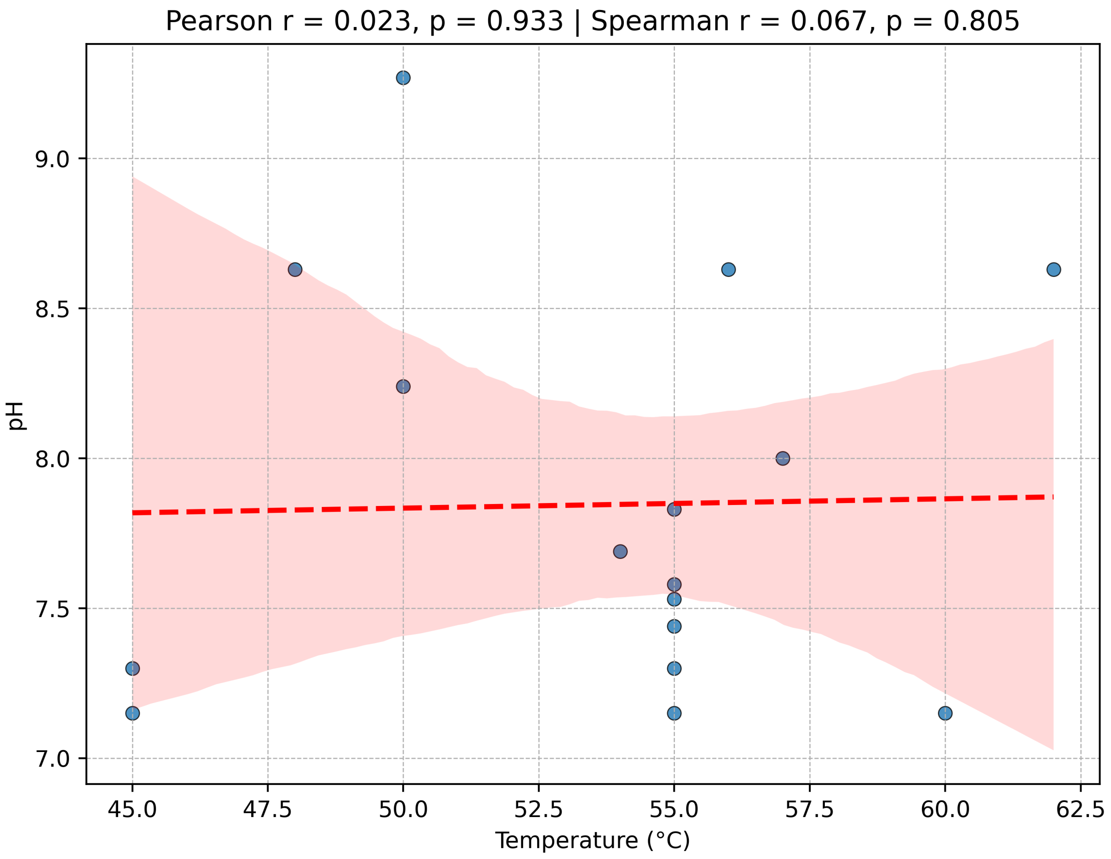


Fig. S3. Correlation between pH and temperature in El Tatio samples. Both variables show statistical independence (Pearson's and Spearman's p), being the explained variability low in both tests. We can safely attribute differences in the explanatory power over the number of sequences per biochemical function (inferred by KEGG orthology) to each variable – an increase in temperature does not influence the observed pH of the samples in our dataset.

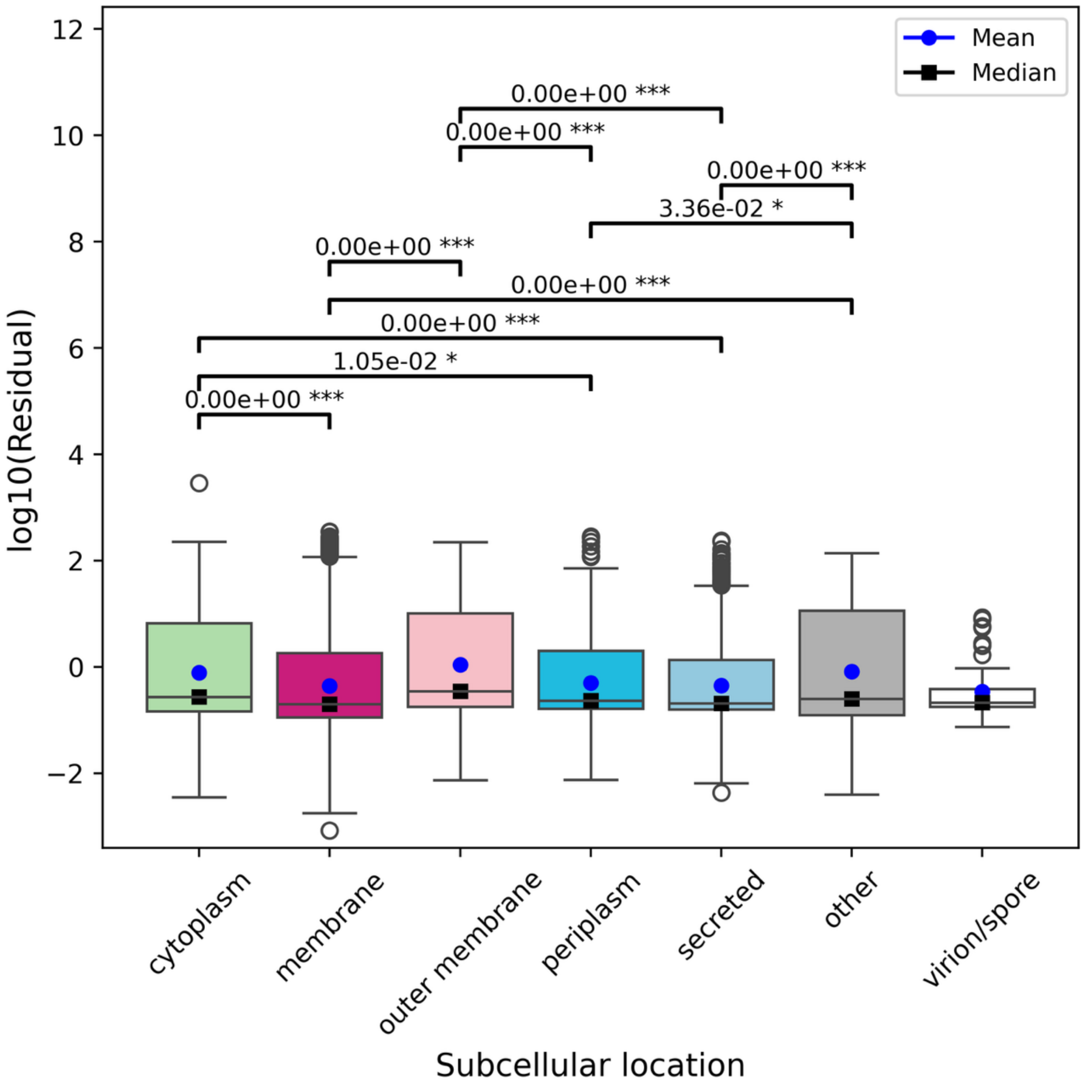


Fig. S4. Abundance-controlled distributions of the by-function sequence diversity per compartment. Pairwise t-tests with Bonferroni correction are displayed when significant. Cytoplasm has higher residuals than membrane proteins, which is consistent with the confined evolution hypothesis. “Membrane” corresponds to any membrane that is not “outer”.

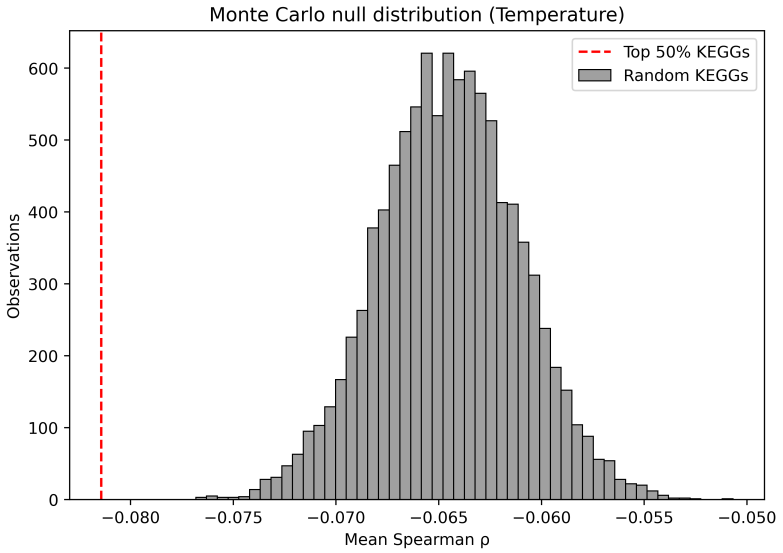


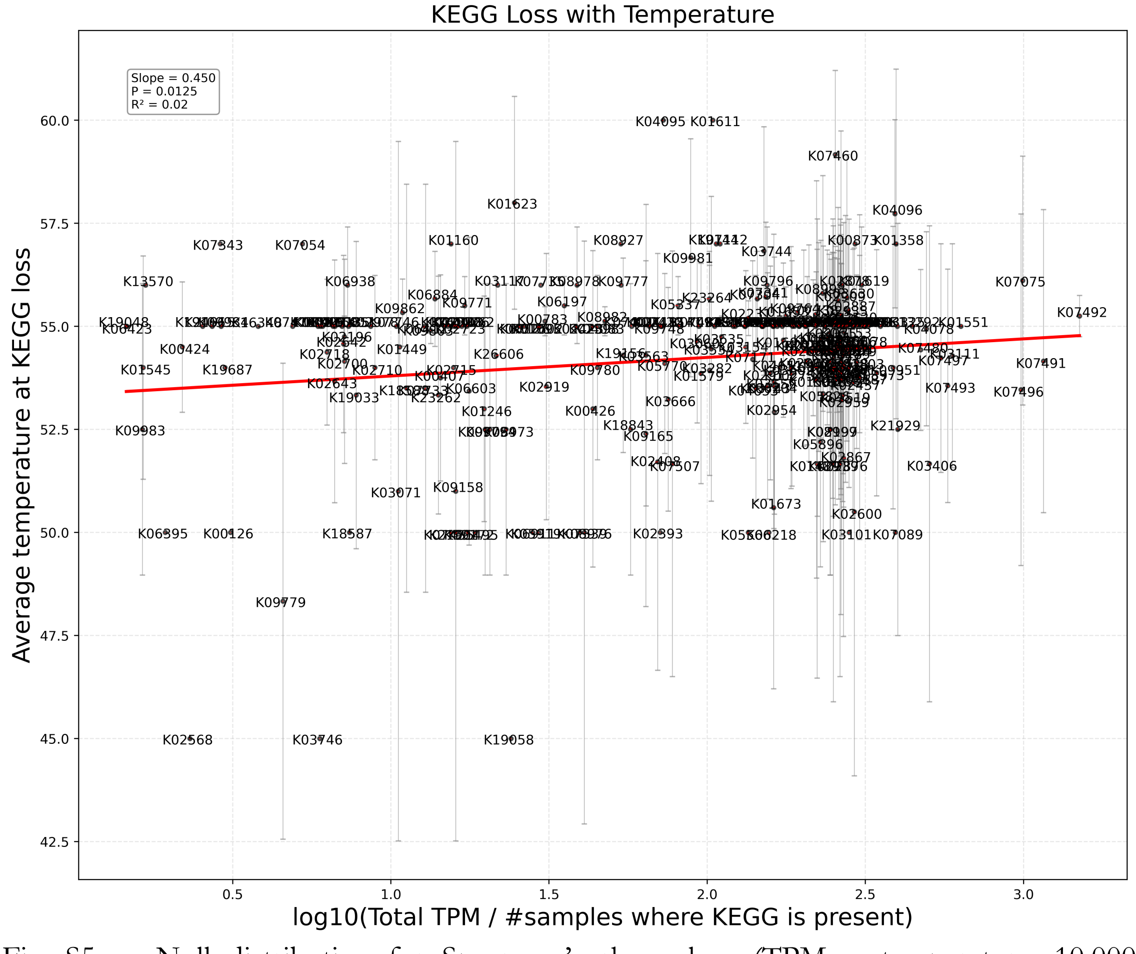


Fig. S5. a. Null distribution for Spearman's rho values (TPM ~ temperature; 10,000 extractions). Monte Carlo test for temperature correlation (Top 50% abundant KEGGs): observed mean ϱ = -0.08; null distribution mean = -0.06 ± 0.003; empirical p-value < 0.001. b. For each KEGG identifier, total abundance was calculated as the sum of TPM values across all taxa and log-transformed. Each point represents one KEGG function and shows its total abundance across metagenomes against the mean temperature of metagenomes in which it was not detected (KEGG is lost). The slope is significantly positive, indicating that the more abundant KEGGs are prone to be absent in any taxa when temperature is higher.

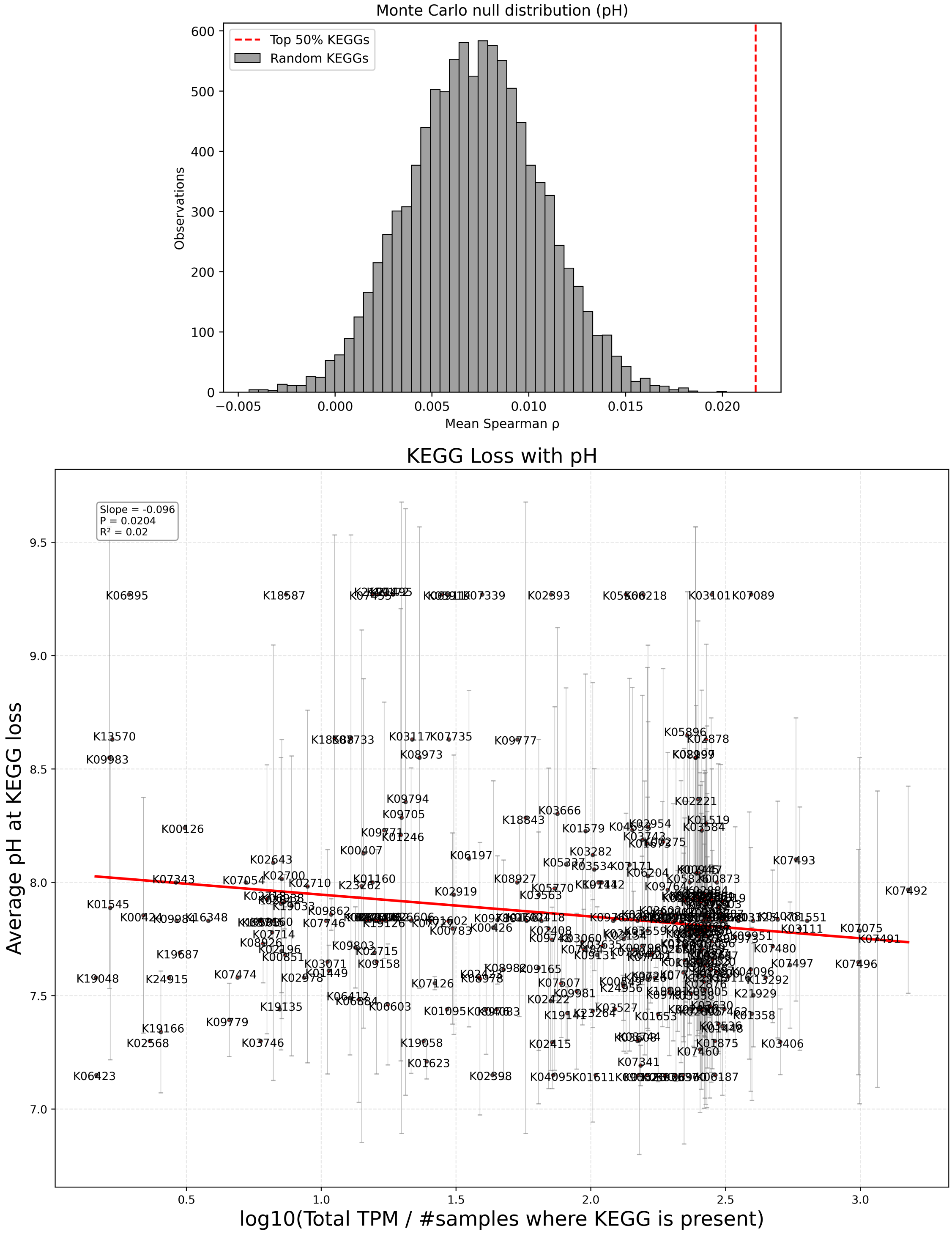


Fig. S6. a. Monte Carlo test for pH (Top 50% abundant KEGGs): observed mean $\varrho = 0.02$; null distribution mean = 0.01 ± 0.003; empirical p-value < 0.001. b. Regressions showing the abundance of KEGGs in average pH conditions where they were absent. The slope is significantly negative, indicating that the more abundant KEGGs are more prone to be absent when pH is lower.

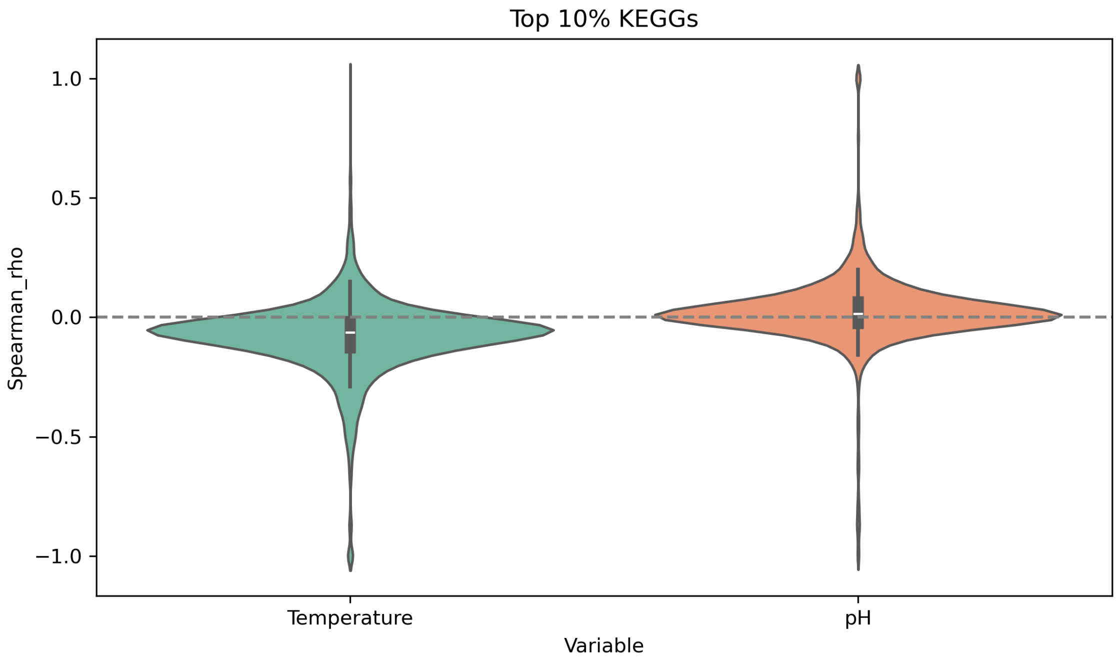


Fig. S7. Comparison of temperature vs pH effects on abundance in Top 10% abundant KEGGs (Spearman ϱ): Mean ϱ for temperature = –0.08 ± 0.15; for pH = 0.02 ± 0.14; t = –28.14, $p < 0.001$.

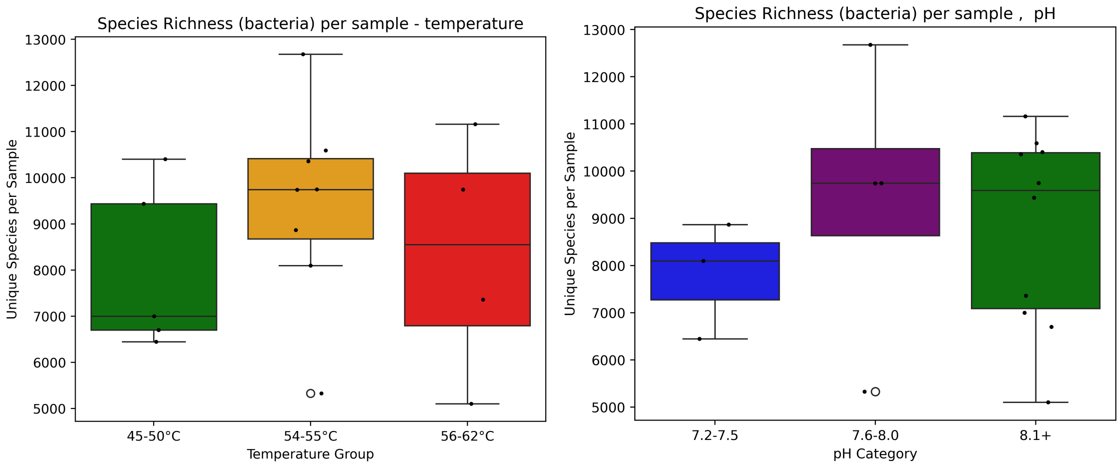


Fig. S8. Bacterial species richness per arbitrary environmental category. None is significantly different. Pairwise t-tests for bacterial richness across pH and temperature categories: pH – 7.2–7.5 vs 7.6–8.0 ($p = 0.40$), 7.2–7.5 vs 8.1+ ($p = 0.35$), 7.6–8.0 vs 8.1+ ($p = 0.74$); Temperature – 45–50°C vs 54–55°C ($p = 0.23$), 45–50°C vs 56–62°C ($p = 0.83$), 54–55°C vs 56–62°C ($p = 0.51$).
